# Validating regional insect pest phenology forecasts using published developmental data

**DOI:** 10.64898/2026.11.13.766408

**Authors:** Darío San-Segundo Molina, Constanti Stefanescu, Ignacio Morales-Castilla

**Author notes:** Correspondence*: Darío San Segundo Molina.

## Abstract

**BACKGROUND:** Crop losses caused by insect pests are expected to increase under global warming, highlighting the need for accurate phenology forecasts to support integrated pest management (IPM). Arthropod pests, as ectotherms, exhibit temperature-dependent development and phenology that can be described by thermal performance curves (TPCs). For decades, developmental models have predicted pest emergence through thermal accumulation approaches, yet it remains unclear to what extent experimentally derived development models can accurately predict field phenology across spatial and temporal scales. Here, we evaluate whether developmental TPCs and linear degree-day models fitted from published experimental data can predict large-scale pest phenology under a rate-summation approach.

**RESULTS:** We compiled published developmental data for the widespread crucifer pest *Pieris rapae* and fitted several TPCs and linear degree-day models. Model predictions were validated against independent long-term monitoring data from the Catalan Butterfly Monitoring Scheme for mainland Spain and the Balearic Islands. Published experimental data accurately predicted large-scale pest emergence, with prediction errors generally below one week. However, predictive performance depended strongly on the spatial and temporal scale considered, with model rankings changing according to the scale and temperature resolution used.

**CONCLUSION:** Published experimental thermal biology data can accurately predict regional insect pest phenology when developmental models are selected according to the intended forecasting scale. These findings provide practical guidance for improving pest alert systems and other decision-support tools, while demonstrating how decades of experimental research can be translated into operational phenology forecasting.

## 1. Introduction

Recent projections of global warming scenarios estimate increased crop damage for nearly half of major crop pests with overall mixed responses – i.e., “winners” and “losers” ^1^ – and average yield losses of 10-25% per degree of warming ^2^. In this context, forecasting the seasonal occurrence and performance of crop pests emerges as an urgent demand to better anticipate impacts and adapt agricultural management practices accordingly. The development rates of arthropod crop pests vary across temperatures predictably following thermal performance curves (*TPCs*, hereafter). The temperature dependence of developmental rates, governed by enzymatic kinetics ^3^, underlies the thermal dependence of population growth rates often linked to fitness ^4^, serving as the basis to inform crop pest warming-induced projections (e.g., Deutsch et al. 2018). Modelling the thermal dependence of development rate makes it possible to estimate the timing of seasonal emergence patterns in crop pest populations, i.e., their phenology (Scranton and Amarasekare 2017). Phenological sensitivity to temperature is expected to cause phenological shifts of insects in response to warming ^7^. Insect crop pests have indeed shifted their phenology in recent decades (e.g., Cohen et al., 2018; Stefanescu et al., 2003), for example advancing their seasonal emergence ^12^ and increasing their voltinism –i.e., the number of generations they undergo within a year ^13^. For multivoltine insects –the case of most important crop pests– phenological advancements enable the completion of additional generations within a year, potentially enhancing their fitness ^14^. In other words, these species would reproduce more frequently within the same year, leading to increased negative effects on plants (Altermatt, 2010; Forrest, 2016).

Process-based models have been amply used to predict phenology in agronomic contexts for decades ^17^, from linear degree-day models ^18^ to Thermal Performance Curves (TPCs, hereafter). Maps of potential voltinism based on thermal accumulation have been extensively utilized in pest risk assessments, especially for invasive crop pests ^19,20^. Phenological forecasts based on modelling how the development rates respond to varying temperatures may also help anticipate the timing and location of potential pest outbreaks, aiding pest management strategies ^21^. The most sophisticated modelling exercises (e.g., Damos et al., 2023; Ponti et al., 2014) are still far from being systematically applied for most crop pests in open, user-friendly tools for growers and policymakers at medium-to-large spatial scales, likely due to their large demand on experimental data. So far, scientifically informed pest management practices have mostly relied on more straightforward approaches based on linear models, taking advantage that they require fewer laboratory data to be parameterized or even using field data ^24^. For instance, the US National Phenology Network (https://www.usanpn.org/data/maps/forecasts) provides a valuable tool based on degree-day models with field validations to elaborate real-time spatio-temporal forecasts of pest occurrence to guide target applications for Integrated Pest Management practices. Similarly, the Degree-Days, Risk, and Phenological event mapping (DDRP) consists of a decision support tool that combines degree-day models and known thermal limits for survival to predict the seasonal occurrence and climatic suitability of several crop pests ^25,26^.

Recently, open-source software has been developed to facilitate fitting nonlinear regression models —e.g., *rTPC* and *nls.multstart* ^27^, *devRate* ^28^ and *mappestRisk* ^29^. Along with the increasingly available experimental data from literature (e.g., synthesized in datasheets of international organizations such as CABI and EPPO), recent advances in the field of thermal biology and ecology may contribute to easily fitting thermal performance curves. Having only slightly higher data demands than linear models, TPCs for development capture the nonlinearity of developmental responses to temperature, thereby achieving more accurate predictions especially under fluctuating thermal regimes and temperature extremes that populations are likely to experience under field conditions ^30,31^. Researchers have increasingly advocated for these TPCs to address intrinsic nonlinearity of the thermal dependence of development rates, since they allow to estimate upper temperature thresholds for development ^32^, to better capture low development at cold temperatures ^31^ and to account for unequal changes in performance with varying temperatures below and above a thermal maxima for development (*T*_opt_), a phenomenon called Jensen’s inequality ^33^.

The effects of temperature fluctuations and extremes on phenology are more likely to be captured at finer spatial and temporal scales. For example, using hourly temperature data can better capture the thermal regimes affecting the phenology of a certain population within the precise location it inhabits. Since temperature has been shown to be the main driver affecting phenology of insects at large scales ^34^, these effects are expected to blur at coarse scales, so that average temperatures are likely to drive the emergence of a phenological signal at macroscales, both spatially (i.e., pooled phenology data of populations across localities) and temporally (i.e., using thermal regimes with daily resolution rather than hourly).

Consequently, we can make several predictions about the scale-dependency of phenological modelling: (1) simpler models, such as linear degree-days, are more likely to accurately predict phenology at coarser spatial and temporal scales, where the effects of temperature fluctuations and extremes is less determinant than at fine scales; (2) that increasing biological realism—by using nonlinear models—and ecological realism—by using hourly temperature data as thermal regimes—can improve the accuracy of phenological predictions, especially at local scales; and (3) that, with careful considerations on thermal biology and ecology of the target species, using disparate source of biological data on development across temperatures can lead to consistent phenological predictions at large spatial scales, but not at local scales.

Altogether, here we aim to address the following questions: (1) to what extent modelling development rates using experimental data leveraged from the crop pest literature can predict observed temporal patterns in pest emergence across spatial and temporal scales? and (2) in which ecological and applied contexts both linear degree-day and nonlinear TPCs enable reasonably accurate phenological predictions for pest management? We address these questions using the well-studied pest *Pieris rapae* as a case study, taking advantage of the availability of both experimental and observational data that enable fitting models and independent validation of their predictions. Addressing these questions should shed light on the existence of middle grounds in the trade-offs between model complexity, data availability and practical application for managers and policy-makers. Successful predictions within this balance would open a door to systematically integrating existing experimental knowledge into inferential schemes.

## 2. Material and Methods

### 2.1 Study system

Here we explore these questions using the small white, *Pieris rapae* L., an abundant, widespread multivoltine butterfly with high migratory rates. This generalist species with winter pupal diapause can attack multiple brassica crops. For species whose biology and thermal ecology has been extensively studied such as *Pieris rapae*, it should be possible to build models and spatial-temporal forecasts for validation against available observational data (e.g., Gordo and Sanz 2006, Colom et al. 2021).

The emergence date (on average) of the first generation of butterfly adults exiting winter diapause is optimal to evaluate the predictions from TPC –phenological – models against observational phenology data. Using this phenophase has three main advantages. For insects showing a partial migration strategy, where a proportion of a population migrates while other individuals remain resident ^37^, as is the case of *P. rapae* populations in our study area ^38^, the emergence of the first generation at a site is primarily driven by local thermal accumulation rather than the arrival of immigrants. In other words, the earliest observed flights likely come from individuals that survived the winter at the site and resumed development with rising spring temperatures after terminating winter diapause, as migration does not occur that early in the season. In contrast, later observations may be biased by the arrival of populations from North Africa and South Spain in late-spring and from Central Europe in autumn, leading to a population with mixed phenological histories. Hence, the occurrence of these first imagoes of the year may reflect the population phenological sensitivity to temperature accumulated in winter and spring after pupal diapause termination ^39^. This thermal accumulation for exiting diapause is likely to follow a typically left-skewed TPC, as documented for its congener *P. napi* ^40^. Second, as a thermophilic species, *P. rapae* can reach short development times at high temperatures (Vives-Ingla et al., 2023; von Schmalensee et al., 2023) allowing short development times at summer. Along with its large intrapopulation variability and migratory ability, this leads to observed overlapping generations of *P. rapae* during summer in Mediterranean areas ^36^. Additionally, although there is no evidence for facultative aestivation of *P. rapae* in Southern Europe, we assume it is likely to occur based on late-summer population dynamics on several recording transects in Northeastern Spain (Appendix S1, Fig. S1–S3), as it has already been documented for *P. brassicae* (Spieth, 2002). These three factors (especially the first two, given the uncertainty regarding the summer diapause of *P. rapae*) make abundance data of first adults in spring the best candidate to examine the accuracy of phenological predictions based on TPC– rate summation approaches.

### 2.2 Development rate data

We retrieved experimental laboratory data on the variation of development days for populations of *P. rapae* across temperatures from two published sources. First, the Gilbert (1988) dataset contains experimental data on populations from three temperate locations with oceanic climate (Canberra in Australia, Vancouver in Canada, and Ascot in the UK) and three Mediterranean locations (Cordoba in Spain, Pompei in Italy and Montpellier in France), at which *P. rapae* has overwinters as a diapausing pupa ^44,45^. This dataset comprises *n* = 32 experimental records of development rates as a response to temperatures in the range 14 – 31°C, which were obtained using WebPlotDigitizer v4.8 (https://apps.automeris.io/wpd4/) from Fig. 1 in Gilbert (1988). Second, the von Schmalensee et al., (2023) experimental dataset comes from Swedish populations, where *P. rapae* is sympatric with *P. napi*. This dataset comprises *n* = 133 experimental records of development rates as a response to temperatures in the range 10 – 35°C. Merging data from populations of different geographic origin to calibrate models is a common practice in the species distribution modelling literature, given it allows a more complete characterization of the species niche and more accurate predictions even if the model is only projected to a limited portion of the species distribution (e.g., Chevalier et al., 2022). We follow a similar rationale here, but test whether making phenological predictions joining largely divergent multi-population data on thermal performance ^47^ leads to higher prediction accuracy (see *Sensitivity analyses* section).

**Fig. 1.**
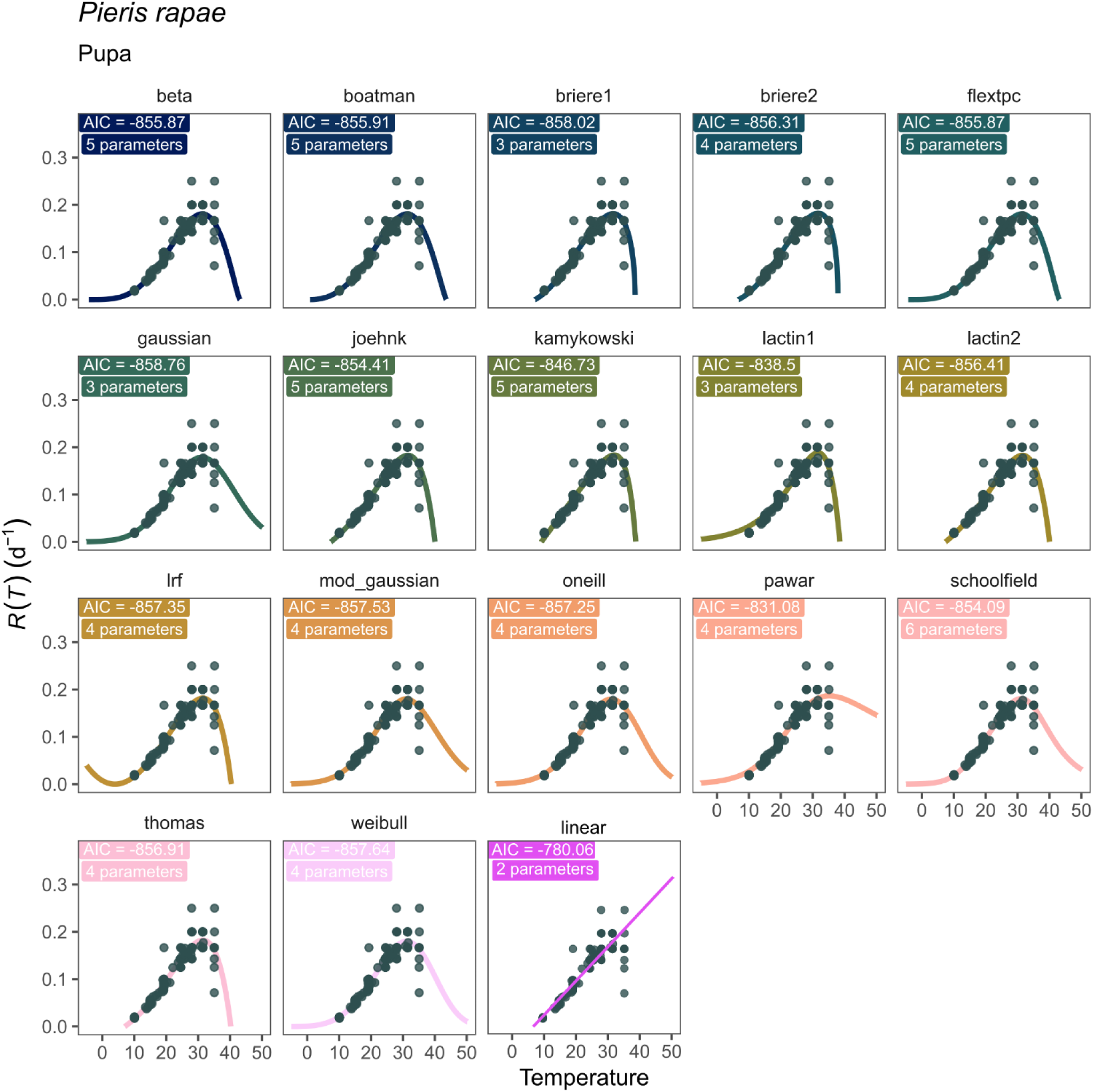
Thermal Performance Curves and linear model fitted for development data retrieved from Gilbert (1988) and von Schmalensee et al. (2023) data sets, as represented by gray points. Each panel includes the model AIC and the number of parameters of the model equation. Panels are ordered alphabetically, except for the linear model, placed last. The *y-*axis represents the development rate (in inverse days) and the *x-*axis represents the temperatures

### 2.3 Observational phenology data of Pieris rapae

In this study, we used observational, long-term data on *Pieris rapae* phenology from the Catalan Butterfly Monitoring Scheme (CBMS hereafter, https://www.catalanbms.org/). This data has been collected across Andorra, Catalonia and Balearic Islands along fixed transects for up to thirty-three years at some locations. At each butterfly transect, counts were repeated weekly, starting from 1^st^ March until 26^th^ September (i.e., a total of 30 recording weeks) whenever minimum meteorological requirements for butterfly activity were met. To approximate the first observed flights of the year based on this abundance data, we discarded locations whose yearly data was >50% absences, since we assumed that the populations may not be established there and the – usually late – observations might be the result of migrations. After these filtering steps, a total of 156 unique transects with altitudes ranging from 0 to 1,720 m (median: 221.5 m) were used for validation (Appendix S1, Fig. S1). We then excluded years with sampling biases (e.g., a likely lack of sampling due to COVID-19 restrictions, Appendix S1, Fig. S2, Fig. S3). Next, the first observations (i.e., at least one individual) of each year for each transect were chosen for validation purposes.

### 2.4 Climate data

We extracted climate data from *EOBS* (https://surfobs.climate.copernicus.eu/dataaccess/access_eobs.php), a 0.1° (∼11 x 11 km at the equator) gridded climate data set with daily minimum, maximum and average temperatures to each transect coordinates from the CBMS data. We then averaged temperatures across transects using *terra* package ^48^ to obtain three single daily temperature values (minimum, maximum and average) for the complete region. The thermal regimes obtained with this extraction are described in Appendix S3 (Fig. S4).

### 2.5 Model fitting, prediction and independent validation

We fitted 18 TPC nonlinear models and one linear degree-day model in R ^49^ using *mappestRisk* package ^29^ along with *devRate* ^28^, *rTPC* and *nls.multstart* ^27^. We included AICs as a metric of statistical fit to compare regression models ^50^.

We predicted phenology of post-diapausing *P. rapae* populations using a rate-summation approach using daily extracted temperatures (minimum, maximum and average) and the development rate models fitted before. For linear, degree-day models we computed rate summation following three different methods: rectangle, or classic degree-day model (based on daily average temperature), triangle and sine (simulating fluctuations between minimum and maximum daily temperatures), following Mirhosseini et al., (2017). For nonlinear, TPC models, rate summation was computed by iteratively summing the rate of development given at a certain temperature over discrete time increments ^31,51^ (Appendix S2). To examine the effect of the downscaling the temporal resolution of temperatures on phenological accuracy of models, we simulated hourly temperatures across the year at 40°N latitude with the *make_hourly_temps* function from the *chillR* package, which models within-day temperature variation using a sinusoidal-exponential approach that incorporates sunrise and sunset times. These hourly temperatures were then used as input to iteratively predict phenology through the rate summation operations. We performed these simulations using both the extracted temperatures at each transect and those averaged across transects.

For validation purposes to examine the predictive accuracy of models, we compared predicted emergences under rate summation against the observed first flights (see above) from the CBMS data, both for each transect and for the pooled data across transects. For this purpose, we used the Root Mean Square Error of Prediction –RMSEP (Appendix S2), whose value indicates the number of days by which predictions and observations differ.

### 2.6 Sensitivity analyses

We conducted two sensitivity analyses to explore whether our results in predictive accuracy of phenological models across scales were consistent to (1) using disparate sources of phenological, observational data for validation; and (2) using disparate sources of developmental data across temperatures, i.e., using development data for populations differing in their geographic origin. For the first sensitivity analysis, we repeated the complete workflow defined here using a validation dataset from Gordo & Sanz, (2006), which compiles first emergences of *P. rapae* across mainland Spain for more than fifty years. For the second sensitivity analysis, we repeated our analytic approach (i.e., curve fitting, making spatial-temporal predictions, comparing predicted values against independent field observations) but fitting the curves using separately the two experimental development data sets (Gilbert 1988; von Schmalensee et al. 2023).

## 3. Results

Fitted models showed small differences in AIC scores, but they largely differed among model equations (Appendix S3, Table S1), especially at the hot- and cold ends of the curves (Fig. 1). The differences in the cold end of the curve are particularly relevant for predicting the phenology of first adult emergence, since the temperatures in the CBMS sites in this season (from 1^st^ of January to first documented flights) lie frequently in this thermal range, particularly the minimum temperatures (Fig. 2). Thus, the estimated cold-developmental threshold slightly differed between most models, with some exceptions (Appendix S4, Table S2).

**Fig. 2.**
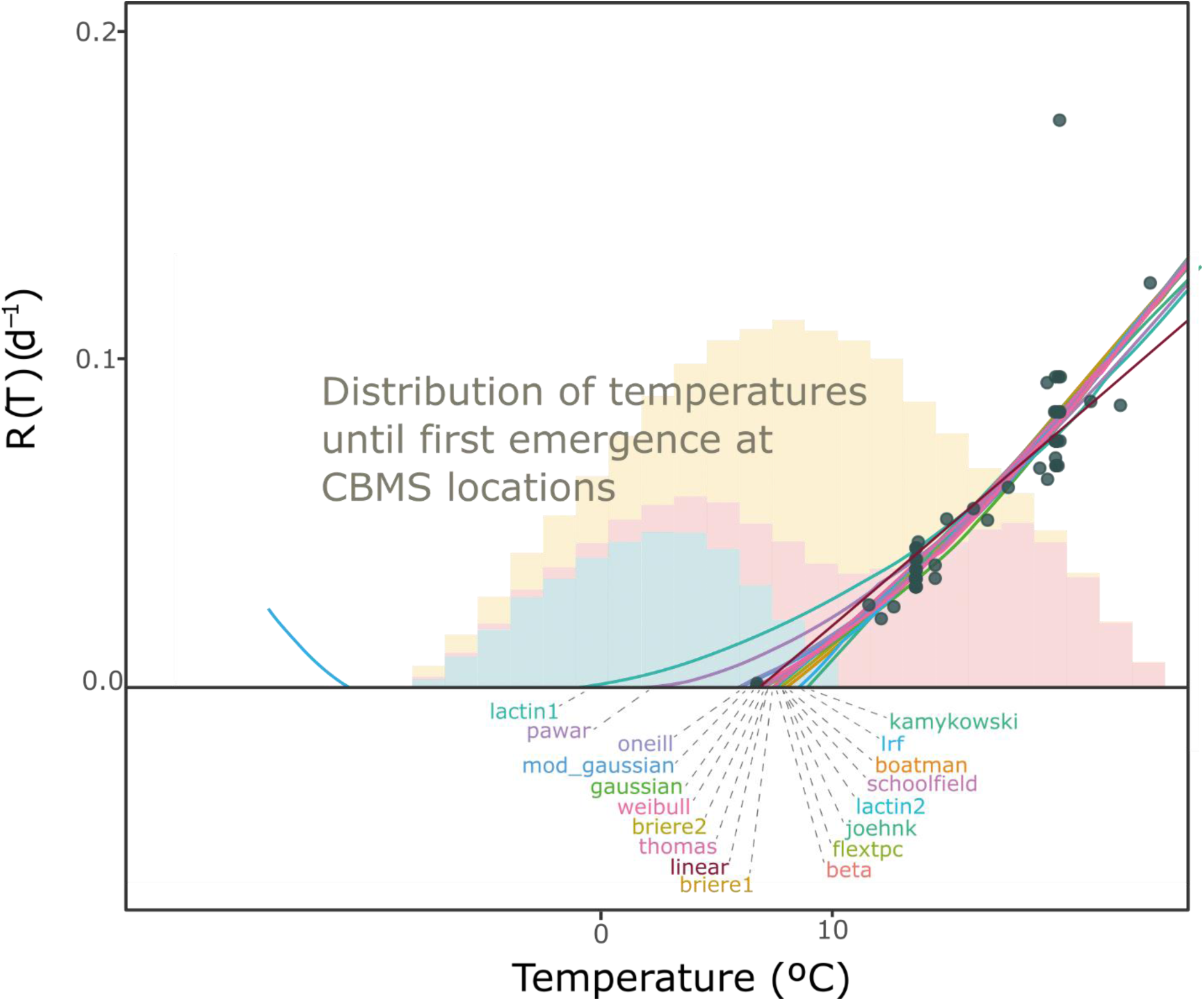
Cold ends of the fitted development models. Points represent the development data retrieved from Gilbert (1988) and von Schmalensee et al. (2023) data sets. The *y-*axis represents the development rate (inverse days), and the *x-*axis represents the temperatures. Each model equation is depicted with a color, as indicated by the labels. These labels are placed according to their cold-end value (Appendix S4, Table S2). The histograms in the background represent the frequency of each temperature (in a sequence of 1°C) in the CBMS locations from 1^st^ January to the average first observed flight that year across transects, for daily minimum (blue), average (yellow) and maximum (red) temperatures. Note that depicted curves and endpoints are an approximation for facilitating visualization with illustrative purposes; values of cold development thresholds and accurate model predictions are given in Appendix S4, Table S2 and in Fig. 1, respectively.

Predictions of adult emergence phenology derived from fitted developmental rate models were generally accurate across both long-term time series and spatially aggregated datasets, with mean absolute errors typically below ±8 days in best fitting models (Fig. 3, Appendix S5, Tables S3– 4). These levels of accuracy were consistent regardless of whether the experimental datasets used to fit the models were considered separately (Appendix S6, Fig.s S5–6). Predictions were also similarly accurate when using an independent data set of field observations for validation (Gordo & Sanz,2006; Appendix S7, Fig. S7). Nevertheless, the models achieving the highest predictive accuracy varied depending on the combination of fitting and validation datasets, reflecting differences among equations and parameterizations of the thermal performance curves and linear development-rate models (Fig. 1, Appendix S3, Table S1).

**Fig. 3.**
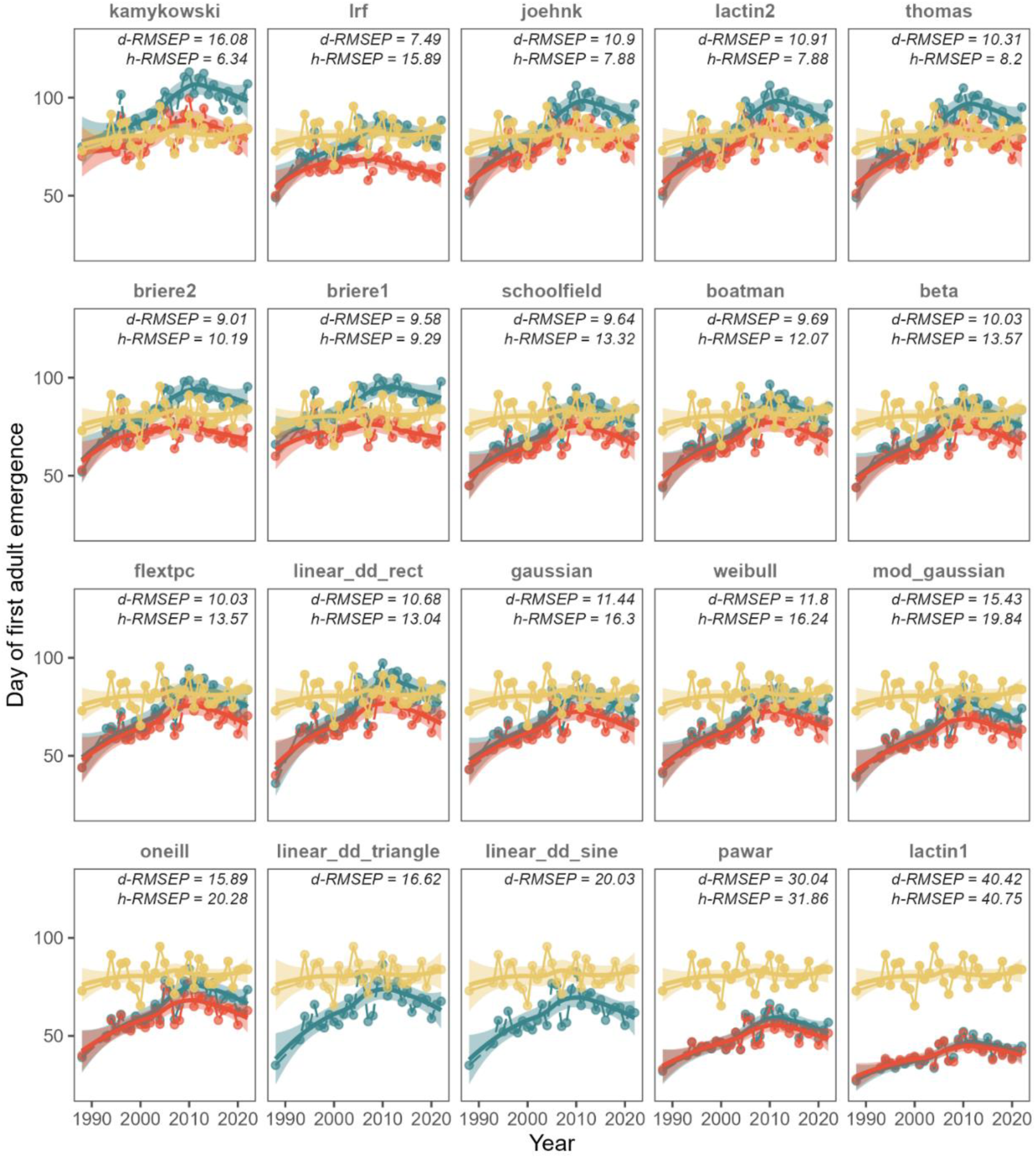
Validation of phenological predictions from TPC and linear models fitting to observational data at CBMS locations. Blue and red points represent phenological predictions from fitted models of first adult emergences, averaged across all transects, using daily and hourly temperature data, respectively. Predictions were accomplished by rate-summation for nonlinear TPC models and by degree-day calculations with rectangle, triangle and sine-methods for the linear model. Yellow points represent the averaged first observations of *P. rapae* adults across CBMS transects. Lines represent smooth trends with a *loess* model intended for facilitating visual comparison. RMSEPs are printed with abbreviations for both daily and hourly temperatures. Panels are ordered based on the lowest value of any RMSEP.

The combined experimental data used to fit the models showed an almost linear increase of development rate with temperature, up to a threshold temperature beyond which development rates decrease sharply (Fig. 1). Several linear and non-linear models did similarly well at fitting curves closely matching these data. For example, nonlinear models *briere1*, *gaussian*, *mod_gaussian, oneill*, *thomas*, *weibull* and *lrf* and have similar statistical performance according to AIC values (ΔAIC < 2) (see Fig. 1), but they slightly differed in their estimated cold developmental thresholds (Fig. 2).

Model accuracy in predicting emergence dates for independent observations (i.e., data by the CBMS) differed substantially among models (Fig. 3), and the rank order of predictive accuracy did not match the AIC-based ranking (Appendix S5, Tables S3–S4).

At the daily temperature resolution, the most accurate models, with errors ranging from 7.4 to 9.7 days, were *lrf, briere2, briere1, schoolfield* and *boatman* (Fig. 3, Appendix S5, Table S3). At the hourly resolution, these models, which ranked from 6.3 to 9.3 days, were *kamykowski*, *joehnk*, *lactin2, thomas* and *briere1* (Fig. 3, Appendix S5, Table S3). Overall, model predictive accuracy worsened when using hourly temperature data by ∼28%, except for the least convex models with higher cold developmental thresholds (Fig. 2), which improved their accuracy at hourly temperatures. Thus, some of the best models overall performed better at the daily time scale (e.g., *lrf* (112%), *gaussian* (42.5%), *schoolfield* (38.2%) and *weibull* (37.6%), Fig. 3, Appendix S5, Table S4).

Using separately the experimental data sets from von Schmalensee et al. (2023) and Gordo & Sanz (2006) to fit models and predicting phenology influenced the predicted temporal patterns of first-adult emergence. TPCs fitted with data from von Schmalensee et al. (2023) often produced phenologies that were earlier than observed, compared with predictions based on TPCs fitted using Gilbert’s (1988) data (Appendix S6, Fig.s S5–6). In other words, the former tended to overestimate development rates—especially when using hourly temperature data—whereas models fitted to Gilbert’s data generally underestimated phenology. However, incorporating hourly temperatures substantially reduced these discrepancies, especially for predictions with Gilbert-only data (Appendix S6, Fig. S6). Yet, several models ranked similarly, with slightly convex TPCs (namely *beta, gaussian, weibull*) ranking consistently among the highest predictive accuracies for the three exercises, i.e., Schmalensee-only (Appendix S6, Fig. S5), Gilbert-only (Appendix S6, Fig. S6) and combined data sets (Fig. 3). Similarly, lactin1 and modified linear, degree-day models (i.e., triangle and sine wave) consistently yielded less precise estimates for the three exercises (Fig. 3, Appendix S6, Fig.s S5–6).

Using a different source of observational data on adult first flights (i.e., the dataset from Gordo & Sanz, (2006)) to validate the predictions for the combined experimental datasets led to similar patterns than the previous sensitivity analysis. In this case, the same slightly convex models at the cold end (e.g., *gaussian*, *beta* and *weibull*) predicted phenology of the first adult generation more accurately than those that had resulted more precise in the CBMS validation exercise (*lrf, briere1, briere2, thomas, joehnk, lactin2)*, whose performance improved for hourly temperature data (Appendix S7, Fig. S7).

Finally, we found higher predictive accuracies (i.e., lower errors at estimating emergences) when we pooled data across transects (Fig. 3, Appendix S5, Table S3) than when we made predictions at the transect level (Appendix S8, Fig. S8).

## 4. Discussion

Our findings confirm that published experimental data on insect thermal performance can support accurate phenological predictions at macroecological scales. However, no single model provides better predictions across all contexts. Instead, model predictive accuracy depends on the ecological conditions of the forecasts, namely the thermal regimes and the spatial and temporal scale at which phenology is modelled.

Using statistical criteria for model selection, such as ranking AICs, R^2^ or sums of residuals, is a standard practice in ecological modelling and forecasting (see e.g., Aghdam et al., 2009), but our results highlight the need to complement the selection process with biological (e.g., the TPC shapes and the diapausing strategies) and ecological (e.g., the spatial and temporal scale and the thermal regimes) information. Here, we model a life stage that requires integrating the TPC across a winter and spring thermal regime near the TPC cold end (Fig. 2). The TPC behavior there determines the prediction success or failure of different equation models but cannot be captured by AICs as source data to fit models lack such cold temperatures.

For example, models with a nearly asymptotic cold-end (e.g., *lactin1, pawar* and *mod_gaussian*) consistently overestimated development when contrasted with the observed populations in the CBMS data, thus predicting emergence before it occurred. Noteworthy, their prediction accuracy was unrelated to their AIC scores (Fig.s 1 and 3, Appendix S3, Table S1). In contrast, when validated against the dataset obtained from Gordo & Sanz, (2006) for mainland Spain, some of the models with lower cold development thresholds, which resulted suboptimal for the CBMS data, emerged then as the best ones at predicting phenology (e.g., *gaussian, weibull, beta, schoolfield, mod_gaussian and oneill*; Appendix S6, Fig. S7). This may occur due to earlier average mean first flights in Spanish Southernmost populations. Alternatively, the discrepancies when validating against both data sets may also arise from methodological differences in their collecting protocols; since we have low detailed information on the protocol of the mainland Spain dataset, we focus our discussion on the CBMS dataset, which has been collected using a standardized, systematic set of protocols.

The predictive accuracy of models with a less convex behavior at their cold end – e.g., *briere’*s, *thomas*, *lrf* and *linear degree-day*’s – is entirely scale-dependent. Thus, which model is more appropriate to predict phenology in an applied pest management context will likely depend on the spatial and temporal scale at which forecasts are conducted. Examining separately the spatial and temporal scale-dependency of phenological forecasts allows drawing some general conclusions.

The **temporal scale** is pivotal to capture the thermal regimes experienced by the organisms, and particularly to incorporate fluctuating temperatures near the cold developmental threshold ^30^. For the daily scale applied to both validation datasets, we found that less convex models at the cold end, usually with higher cold thresholds for development (particularly *kamykowski* for the main validation exercise, see Appendix S4, Table S2) underestimated development at the thermal regimes experienced by the first generation of the butterfly at our sites and propagated these differences throughout the season, thus predicting too late first adult emergences (Fig. 3) at large spatial scales. This tendency to underestimating development was reduced when using the hourly resolution temperature data, improving accuracy (Fig. 3), especially when using data from mainland Spain of Gordo & Sanz (2006), whose populations are presumably less cold-adapted. In this case, several less convex models (e.g., *linear, briere1, briere1, thomas, lrf*) corrected their underestimation of development when using hourly temperature data (Appendix S6, Fig. S7) or, in the case of the linear degree-day model, when applying modifications to compute degree-days at daily scales, such as triangle and sine wave (Fig. 3, Appendix S5, Table S3). These patterns agree with previous findings by Khelifa et al. (2019), who pointed out the difficulty of lab-parameterized TPCs to translate into field data when fluctuating hourly temperatures fall at the cold-end of the TPCs, as rate summation in this context may shift the estimated cold threshold for development to cooler values ^54^.

These findings of the context-dependency of model performance for predicting phenology is analogous to other modeling frameworks, such as species distribution modeling, where different algorithms improve accuracy by emphasizing either true positives or true negatives. In such cases, combining models –typically through averaging– has been shown to reduce predictive uncertainty ^55^. Similar ensemble modeling approaches could be explored for using thermal performance curves to forecast crop pest phenology (e.g., Bates & Bertelsmeier, 2025).

The **spatial scale** also plays a critical role in the accuracy of phenological predictions. Models performed better when data were aggregated across CBMS transects (i.e., when using a coarse spatial scale) than at the local transect level (Fig. 3 against Appendix S8, Fig. S8). Besides the overall decrease in predictive performance, we found that hourly temperature resolutions improved accuracy for suboptimal models (namely the less convex ones, such as *briere1, briere2, kamykowsky, joehnk, lactin2, thomas* and *linear*), while the most consistent models of the aggregated data in Fig. 3 and in the sensitivity analyses (i.e., *gaussian, beta, schoolfield*) still ranked among the top ones here. Pooling sample data across the spatial range of the species allows better characterization of their phenological responses as parametric predictions approach observed values ^57^. As a consequence of the well-known statistical aggregation effects across scales in ecology ^58^, pooling data allows the detection of large scale *phenological responses to temperature*, at the cost of obscuring local scale phenological variability ^59^. Aggregating temperature inputs themselves can introduce a similar bias, since averaging ignores the distribution of temperatures around the mean and can therefore misrepresent performance near thermal optima ^60^. This is consistent with findings showing that, as spatial scale increases, temperature becomes the main predictor of butterfly phenology ^34^. Nonetheless, first emergence data in some CBMS transects – specifically at costal lowlands with mild winters that would enable mid-winter first adult emergencies of *P. rapae* – may reflect arbitrary decisions based on time calendars of monitoring (i.e., first days of March) rather than true first emergences; if that was the case, more convex models might perform better at reflecting true populations’ phenological responses that the data itself might not currently inform. Most notably, these findings on the scale-dependency of predictive performance of phenology models reinforce the usefulness of using development models parameterized with experimental data to make phenological predictions at large scales, thereby informing forecasts for pest management applications at regional scales.

Finally, discrepancies in the model rankings between the main results and the separate analyses of the two development datasets (Appendix S6, Fig.s S5–6) may be caused by intrinsic variation of thermal responses of the source populations. Lower thermal limits for development have been shown to decrease with latitude ^14^. Accordingly, previous findings in Buckley (2022) show that higher latitude populations are predicted to occur earlier at mid latitudes. This may explain why models fitted to the von Schmalensee (2023) data separately, originally collected for Scandinavian populations, result in earlier predictions (Appendix S5, Fig. S5) than those fitted separately to Gilbert’s data (Appendix S5, Fig. S6), originally collected from Mediterranean-to-temperate regions. This finding agrees with previous patterns documented for *Pieris rapae* on local adaptation ^61^ that would lead to phenological adaptations to different biotic and abiotic conditions, as also found for other pierids (Kerr et al., 2020; Siemers et al., 2024; Spieth, 2002).

Our findings must be interpreted with caution and should not be extrapolated to phenological forecasts attempted to target local outbreaks, since higher temperature variability at local scales than at broader scales may propagate predictive errors in development models ^30^, as also observed here. Local adaptation of pierids could further generate alternative mechanisms modulating phenology ^61^. For example, the photoperiod’s effect on diapause induction has been shown to vary across latitudes, i.e., shows local adaptation ^63,64^, and has been shown to affect Iberian butterfly phenology at local scales ^65^. Previous work using CBMS data has found that precipitation delays butterfly phenology ^8^. Moreover, warming and aridification have been linked to population declines in several of these same CBMS communities ^66^, which could reduce the reliability of first-emergence records in the most affected transects through lower detectability rather than genuine phenological shifts All these phenological adaptations are likely to affect the phenological sensitivity to temperature, especially at local scales, leading to lower predictive accuracy of phenological models based solely on thermal dependence of development rates.

Accordingly, future modelling efforts targeting local scale predictions of crop pest phenology, which are particularly relevant for grower decision-making, could be improved in several ways. First, rate summation models can benefit from more biologically realistic temperature measures. Using air temperature as the predictive variable rather than actual body temperatures may bias phenological predictions of seasonal emergences at local scales, since the spatial variability of surface temperatures at which insects’ life stages develop is usually larger than the 1.5m height air temperatures measured by weather stations (Greiser et al., 2022; Kearney et al., 2010). In other words, interpolated macroclimatic data of the kind commonly used in forecasting tools can diverge substantially from the operative temperatures actually experienced by pest organisms in the field (Mosedale et al., 2024). Alternatively, using microclimatic operative temperatures more realistically captures the temperatures experienced by insects, e.g., at leaves surfaces and mesophyll exposed to sun/shade conditions ^41,68^, inside fruits ^69^ or in the soil. This may help to identify when and where thermal buffering opportunities are removed due to the combined effects of warming and aridity on species with cold thermal niches ^41,66^. For this purpose, biophysical models aim to mechanistically approximate the fundamental thermal niche of species based on increasing experimental data and modelling tools ^67,70^ and can be an interesting avenue for modelling studies and applications targeting pest management at local scales. Second, the role of chilling – i.e., cold accumulation – should be further incorporated into more specific modelling approaches for entering and terminating diapauses whenever this knowledge is available, since it has been suggested to accelerate butterflies’ development after termination of pupal diapause ^71^. This diapause termination has been observed to follow a right-skewed TPC maximizing development at cold – but not extremely cold – temperatures for *Pieris napi* ^40^. Third, these events of diapause induction and termination are usually affected by photoperiod interacting with temperature ^15^, and promising studies have recently included it in modelling tools (Grevstad et al. 2022, Damos et al. 2023). And finally, and contrarily to the generalist characterization of *P. rapae,* modelling phenology of specialist insect pests may require accounting for the developmental response of the host plant (e.g., Pulatov et al. 2016, Pollard et al. 2020, Castex et al. 2023), as the emergence of phenological asynchronies due to differential responses to warming may turn into phenological mismatches with fitness cost for the insects ^76^. Indeed, in the same Mediterranean butterfly communities studied here, temperature-driven shifts in phenology and abundance have been shown to reshape species interactions through rewiring ^77^, illustrating how the phenological changes we model can propagate beyond single-species dynamics. Finally, incorporating cohort-based approaches for targeting local outbreaks might improve forecasts (e.g., Marchioro et al., 2016).

Altogether, these findings reinforce that validating the outputs of process-based models with field experimental studies (e.g., Greiser et al. 2022) is desirable, whenever enough data is available, to develop forecasting tools, especially at local scales. Insights from different perspectives can contribute to this objective. For example, development of citizen science projects (see e.g., Mosquito Alert https://www.mosquitoalert.com/ applied to modelling climatic suitability and risk predictions of Asian tiger mosquito, e.g., Pardo-Araujo et al., 2024) and monitoring schemes would help obtain multi-scale observational data. In this sense, large observational data would enable to validate models for the entire phenological cycles rather than single events. Such data would capture more realistically the true phenological responses within the population ^80^. Also, the development of easy-to-use tools for applied purposes with real-time forecasts such as the above-mentioned DDRP ^26^ and the US National Phenology Network, can facilitate producers to continuously submit data on pest outbreaks and emergence patterns that may refine the models with field validation.

In conclusion, our study illustrates the scale dependence of phenological responses to climate, i.e., how spatial scales shape phenological sensitivities ^81^. Gathering sparse experimental and observational data from heterogeneous populations is unable to adequately predict phenology at local scales that might guide *on site* custom pest management practices relevant for growers. However, it can lead to accurate phenological predictions at large spatial scales, which could inform regional pest management policies (e.g., allowing agricultural agencies to issue alerts in advance of outbreaks, to coordinate synchronized sanitation measures or to restrict the temporal windows for pesticide application). In other words, we found that inferring insect phenology at large scales is feasible even with disparate sources of experimental data for the target species under biological and ecological reasoning. This suggests promising opportunities to use largely available physiological data to inform pest management policies at larger scales. These regional scales are usually those at which public institutions make policies and at which pest alert tools and decision support systems are usually developed. More importantly, since several crop pests lack experimental data on development across temperatures, these results reinforce the need to collect new data to enable reasonably accurate phenological forecasts at these geographic scales.

## Notes

https://github.com/dario-ssm/dev_phen_pierisrapae

### Competing Interest Statement

The authors have declared no competing interest.

### Funding

- Ministerio de Ciencia, Innovación y Universidades: FPU20/05528, PID2023-152329OB-I00
- Generalitat de Catalunya
- Barcelona Provincial Council

