## Supplementary Information file for "Validating regional insect pest phenology forecasts using published developmental data"

### Appendix S1. CBMS data:

The subset of the Catalan Butterfly Monitoring Scheme (CBMS) used in our study consisted of 156 transects located in Catalunya, Illes Balears and Andorra Figure S1.

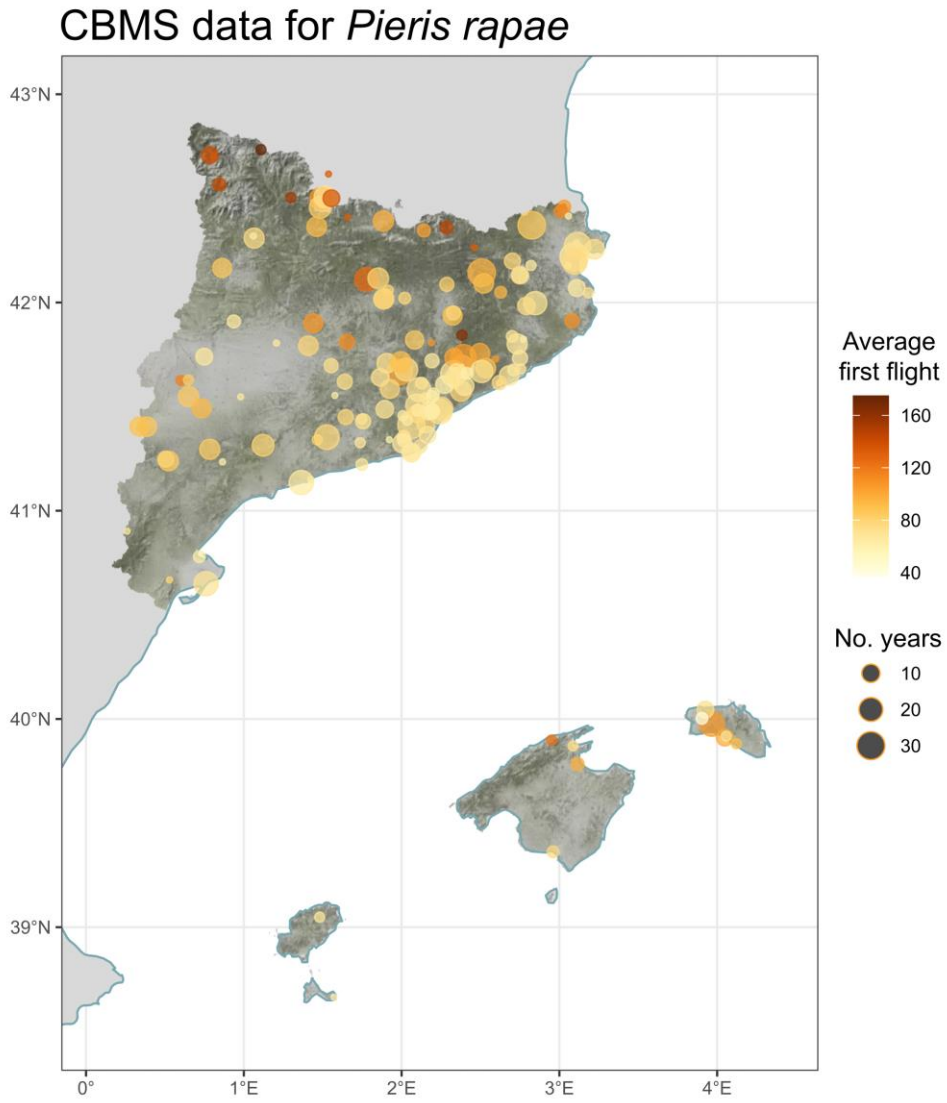


**Figure S1**: Map showing the location of the transects from the CBMS used in our study. The dot size represents the number of years sampled (note that only transects with phenology data for at least ten years were retained). The color of the point represents the average day of year (Julian day) at which the first flight is reported. The tiles represent the LiDAR coverage obtained from *mapSpain* R package *(Hernangómez, 2024)*.

After obtaining these 156 unique transects, we excluded years likely to have sampling biases, such as lack of sampling due to COVID-19 restrictions (see examples for transects 89 and 106 in Figure S2 and Figure S3, respectively).

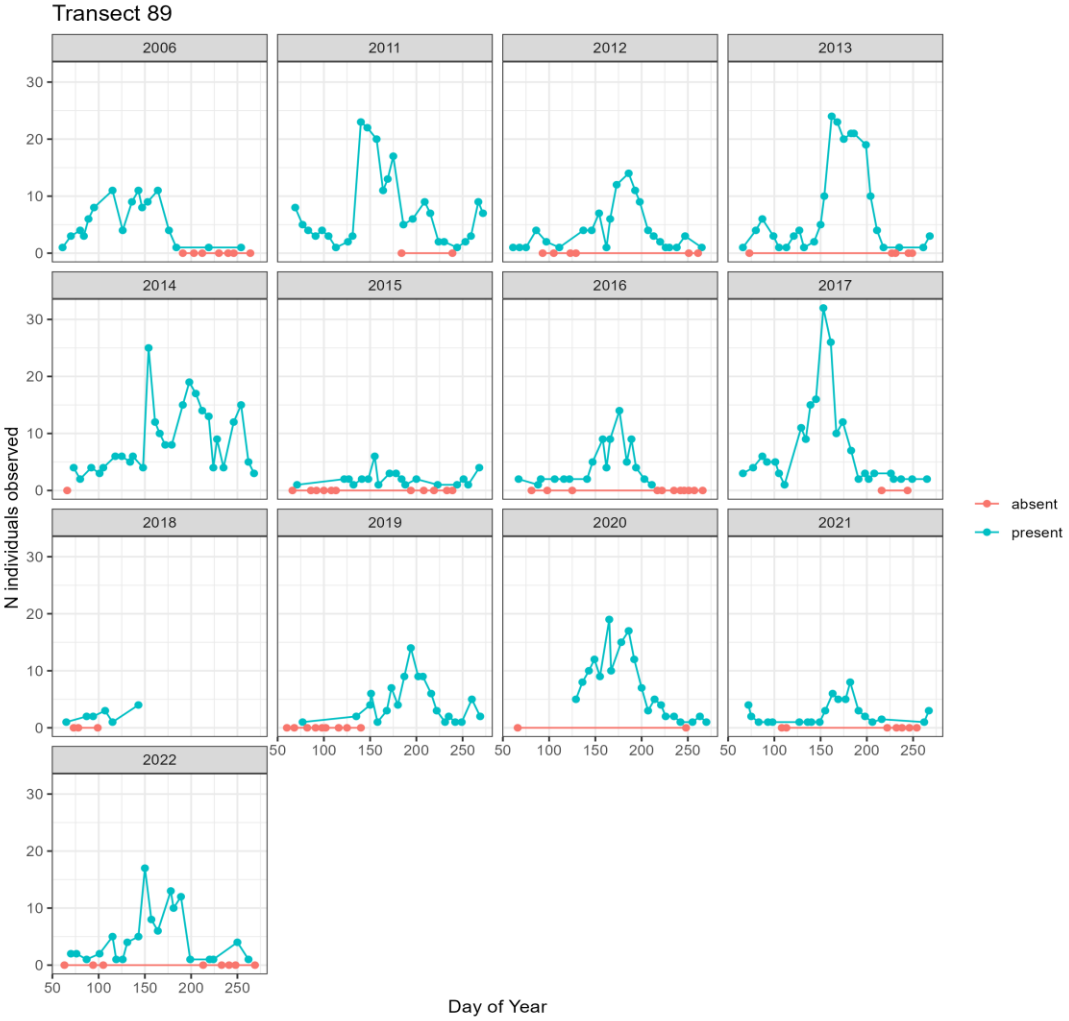


**Figure S2**: Sampling trends in transect 89 of CBMS data across years. Documented absences and presences are represented as red and blue points, respectively. Note that year 2020 lacks both data points for a period between the ~and ~125th days of the year, which coincide with the period of major mobility restrictions for non-essential activities in Spain due to COVID-19.

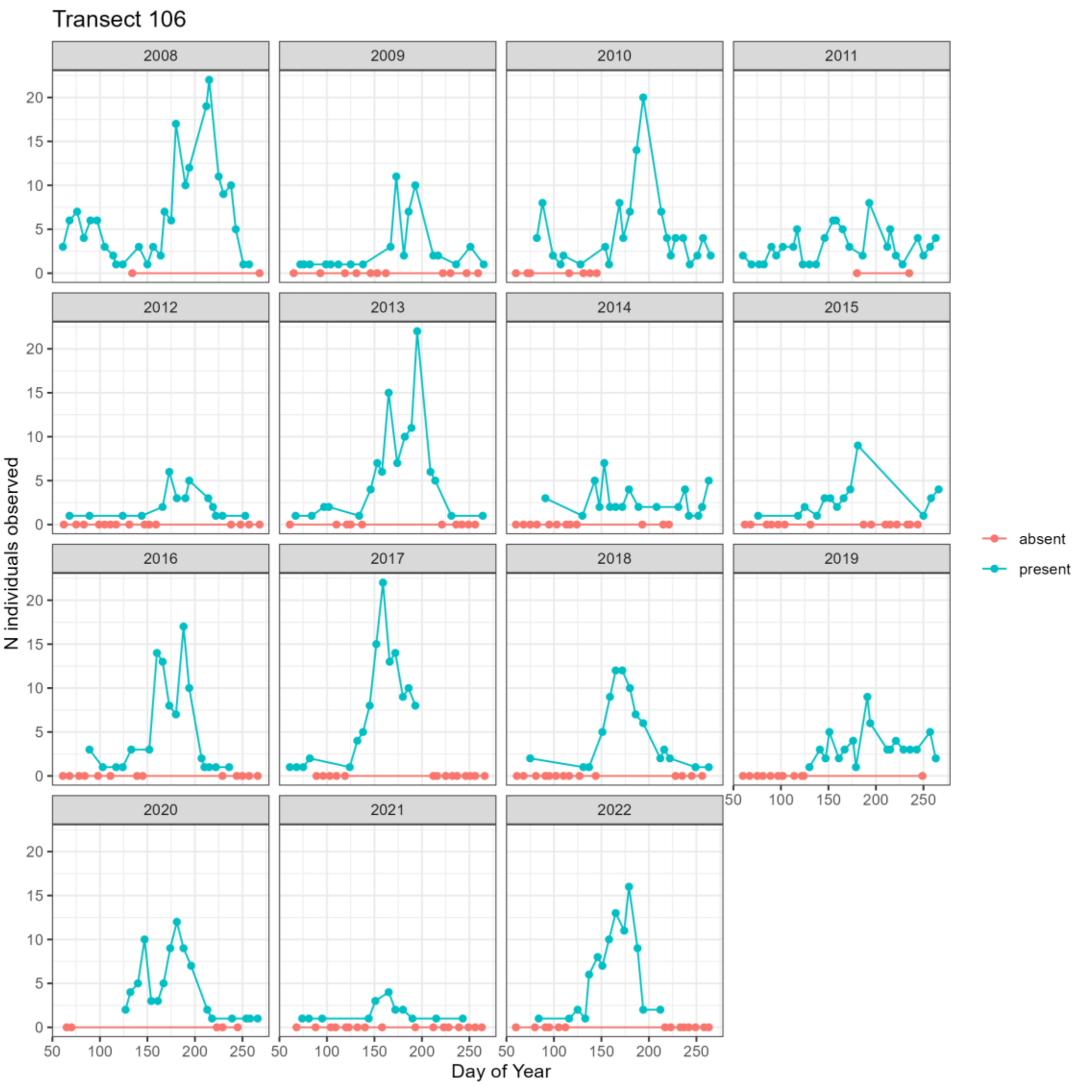


**Figure S3**: Sampling trends in transect 106 of CBMS data across years. Documented absences and presences are represented as red and blue points, respectively. Note that year 2020 lacks both data points for a period between the ~and ~125th days of the year, which coincide with the period of major mobility restrictions for non-essential activities in Spain due to COVID-19. Despite 2019 lacks presence data until a similar date than 2020, absences were reported all the weeks before, as opposite to 2020.

### Appendix S2. Phenological predictions based on rate summation: extended information

#### *a) Rate summation procedure*

For phenological predictions based on nonlinear models, physiological age can be estimated using the rate summation approach as first described by Stinner et al., (1974). In this method, the definite function can be in practice numerically approximated by iteratively summing the rate of development given at a certain temperature over discrete time increments (Liu et al., 1995; Stinner et al., 1974) (Equation 1); for detailed mathematical background see (Curry et al., 1978).

$$\int_{t_0}^{t} R\ \ T\ \ dt \approx \sum_{t_0}^{t} R\ \ T\ \ \Delta t\ \ , \qquad 1$$

where $t$ indicates the instantaneous time (or time increment, $\Delta t$) at which each temperature $T$ is measured. Since $R\ \ T$ represents, by definition, the inverse of the number of days at that temperature required to fulfill development, this value can be divided by the time increment to compute the fraction of development completed during that time interval. For example, a given life stage of an organism for which a daily temperature of 25ºC corresponds to a development rate of $0.2\ \mathrm{days}^{-1}$; in other words, that life stage would need $1/0.2\ =\ 5\ \mathrm{days}$ with a constant temperature of 25ºC to fulfill development and proceed its life cycle. Similarly, if the time interval of interest is 1 hour instead of 1 day, and given the same temperature of 25ºC corresponding to a development rate of $0.2\ \mathrm{days}^{-1}$, after that one hour the life stage will have completed a portion of development corresponding to $\frac{\frac{1}{24}}{\frac{1}{0.2}} = 0.0083$, i.e., a $0.83\ \%$ of development required to proceed to the next life stage. For an illustrative example, we recommend checking the Figure 2 in Greiser et al. (2022).

For linear models, we followed the classic approach of degree-days after Campbell et al., (1974). It can be regarded as a particular case of rate summation. A typical linear model describing development rate variation across temperatures follows:

$$R\ \ T\ \ = a + b.T,$$

$$\text{where: } T_L = -\frac{a}{b}$$
$$\text{and: } K = b^{-1}$$

Here, $T_L$ is the *lower developmental threshold* or *thermal base,* i.e., the temperature value below which no development occurs; and $K$ is the heat constant, i.e., the number of degree-days necessary for the given life stage to fulfill development. In the average method (Moore and Remais, 2014; Sampaio et al., 2021), the degree-days accumulated in a day *i* are calculated as $\delta_i = T_i - T_L$. Hence, the accumulation of degree-days in a given time $t$ can be calculated as follows:

$$k\ t\ = \int_{t_0}^{t} \delta\ \ t\ dt \approx \sum_{t_0}^{t} \delta \cdot \Delta t \qquad 2$$

Then, when the $k\ t$ value obtained in Equation 2 is lower than the parametric heat constant $K$ (i.e., $k\ t\ < K$, the life stage is still undergoing development (or "accumulating heat") until $k\ t\ \geq K$, when it proceeds to the next life stage.

Different methods to estimate daily degree-days across different time increments have been suggested, namely the above-described *daily average*, single and double triangles and single and double sine waves (Allen, 1976; Mirhosseini et al., 2017; Moore and Remais, 2014). In our study, when using simulated hourly temperatures, we simply calculated the degree-hours, i.e., $\delta_{\text{hour}} = T_{\text{hour}} - T_L$ and increased the thermal constant up to $24 \cdot K$.

##### *b) Thermal regimes*

The thermal regimes at which phenological predictions are made are pivotal for ecologically considering the models' performances and their usefulness at each case (Schmalensee et al., 2021; see e.g., Vives-Ingla et al., 2023). We illustrated these thermal regimes across which rate summation is calculated for both daily – from raw climate data – and hourly (using simulations from chillR package (Luedeling et al., 2024)– temperature temporal resolutions, and for both the mainland Spain validation case using Gordo and Sanz, (2006)'s data and for the the CBMS validation case (Figure S7, Figure S8)

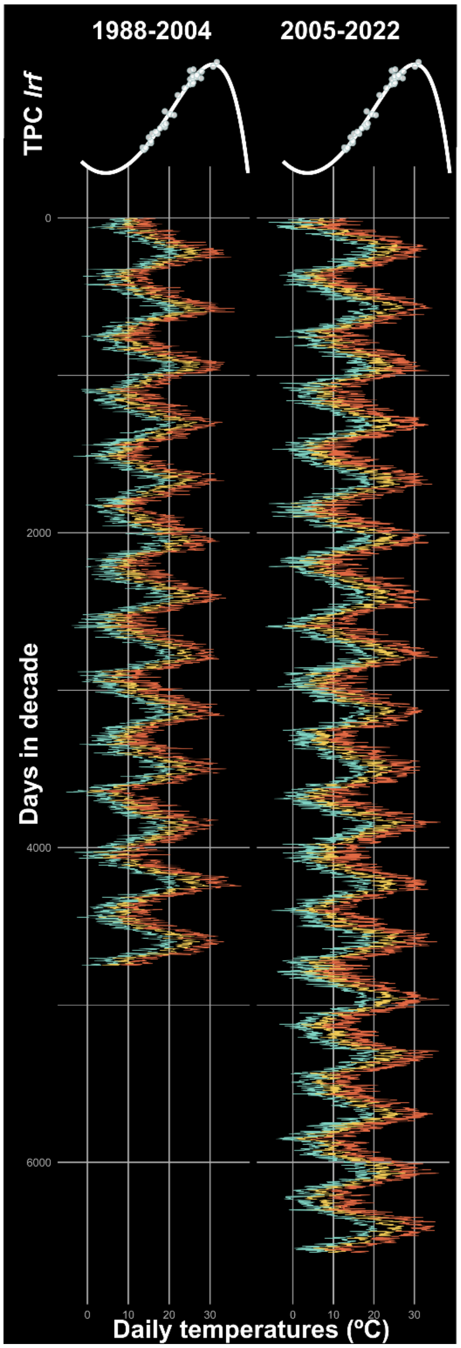


**Figure S4**: Thermal regimes for the Catalunya and Illes Balears case study with validation –observational *P. rapae* phenology data– from the CBMS and climate data from E-OBS (https://surfobs.climate.copernicus.eu/dataaccess/access_eobs.php). For illustrative purposes, we drew the fitted *lrf* TPC above the regimes, as it was discussed as one of the best options (see Discussion in the main text). Red, blue and yellow lines represent the daily maximum, minimum and average temperatures, respectively. Each column in the panel represents one decade and a half.

##### *c) Predictive accuracy of TPC models: RMSEPs*

Root Mean Square Error of Prediction (*RMSEP*) was calculated using *chillR* package (Luedeling et al., 2024) and following Chuine et al., (2016):

$$RMSEP = \sqrt{\frac{1}{N}\sum_{i=1}^{N}\left(N_{PP}\ i\ - N_{PO}\ i\ \right)^2} \quad ,$$

where $RMSEP$ represents the quadratic distance between the observed ($N_{PO}$) and the predicted ($N_{PP}$) phenologies, so that lowest *RMSEP* values indicate higher predictive accuracy.

### Appendix S3: Fitted models (equations, estimated and calculated parameters and source references)

**Table S1.** Models fitted to development rate data across temperatures for the combined data sets from Gilbert, (1988) and von Schmalensee et al., (2023). The table shows the model equations, the source reference where these equations are first described, and the estimated parameters of the models with their corresponding standard error). This last column also includes the Akaike Information Criterion (AIC) score.

| Model name | Equation | Reference | Parameter and AIC (estimate ± se; AIC) |
|---|---|---|---|
| *beta* | $R(T) = \frac{a\left(\frac{T-b+\frac{c\ d-1}{d+e-2}}{c}\right)^{d-1}\cdot\left(1-\frac{T-b+\frac{c\ d-1}{d+e-2}}{c}\right)^{e-1}}{\left(\frac{d-1}{d+e-2}\right)^{d-1}\cdot\left(\frac{e-1}{d+e-2}\right)^{e-1}}$ | (Niehaus et al., 2012) | $a = 0.181 \pm 0.003$<br>$b = 31.3 \pm 0.478$<br>$c = 47.4 \pm 28.20$<br>$d = 4.97 \pm 4.6$<br>$e = 2.32 \pm 2.06$<br>$AIC = -855.87$ |
| *boatman* | $R\ T = r_{max} \cdot (\sin(\pi \left(\frac{T-T_{min}}{T-T_{min}}\right)^a))^b$ | (Boatman et al., 2017) | $r_{max} = 0.181 \pm 0.003$<br>$T_{min} = 0.879 \pm 11.4$<br>$T_{max} = 43.6 \pm 8.27$<br>$a = 2.04 \pm 0.70$<br>$b = 1.14 \pm 1.26$<br>$AIC = -855.91$ |
| *briere1* | $R\ T = aT\ T - T_0\ \ T_L - T^{\frac{1}{2}}$ | (Briere et al., 1999) | $T_0 = 7.22 \pm 0.68$<br>$T_L = 38.6 \pm 0.43$<br>$a = 8.9e-05 \pm 4.9e-06$<br>$AIC = -858.02$ |

| Model name | Equation | Reference | Parameter and AIC (estimate ± se; AIC) |
|---|---|---|---|
| *briere2* | $R\ T\ = aT\ T-T_0\ \ T_L-T^{\frac{1}{b}}$ | (Briere et al., 1999) | $T_0 = 6.76 \pm\ 1.21$<br>$T_L = 37.90 \pm 1.30$<br>$a = 1.03e-04 \pm 2.75e-05$<br>$b = 2.26 \pm 0.56$<br>$AIC\ =\ --856.31$ |
| *flextpc* | $R\ T\ =\ r_{max}\left[\left(\frac{T-T_{min}}{\alpha}\right)^{\alpha}\cdot\left(\frac{T_{max}-T}{1-\alpha}\right)^{1-\alpha}\cdot\left(\frac{1}{T_{max}-T_{min}}\right)\right]^{\frac{\alpha\cdot\ 1-\alpha}{\beta^2}}$ | (Cruz-Loya et al., 2024) | $T_{min} = -4.28 \pm\ 18.4$<br>$T_{max} = 43.1 \pm 10.5$<br>$r_{max} = 0.181 \pm 0.003$<br>$\alpha = 0.750 \pm 0.098$<br>$\beta = 0.188 \pm 0.101$<br>$AIC\ =\ -855.87$ |
| *gaussian* | $R\ T\ =\ r_{max}\cdot e^{(-0.5(\frac{\lvert T-T_{opt}\rvert}{a})^2)}$ | (Lynch and Gabriel, 1987) | $r_{max} = 0.179 \pm 0.024$<br>$T_{opt} = 23.7 \pm 0.9$<br>$a = 10.0 \pm 0.4$<br>$AIC = -858.76$ |
| *joehnk* | $R\ T\ =\ r_{max}\cdot\left(1+a\left(\ b^{T-T_{opt}}-1\ -\frac{\ln b}{\ln c}\cdot\ c^{T-T_{opt}}-1\right)\right)$ | (Jöhnk et al., 2008) | $r_{max} = 0.183 \pm 0.003$<br>$T_{opt} = 31.5 \pm 0.4$<br>$a = 41.3 \pm 80200$<br>$b = 1.11 \pm\ 3.88$<br>$AIC\ =\ -854.41$ |
| *kamykowski* | $R\ T\ = a\cdot(1-e^{-b\cdot\ T-T_{min}}\ )\cdot(1-e^{-c\cdot\ T_{max}-T}\ )$ | (Kamykowski, 1986) | $T_{min} = 9.17 \pm 0.69$<br>$T_{max} = 38.8 \pm 1.42$<br>$a = 15.6 \pm 562$<br>$b = 5.9e-04 \pm\ 2.13e-02$<br>$c = 0.309 \pm\ 0.182$<br>$AIC\ =\ -846.73$ |

| **Model name** | **Equation** | **Reference** | **Parameter and AIC** (estimate ± se; AIC) |
|---|---|---|---|
| *lactin1* | $R\ T\ = e^{\rho T} - e^{\left[\rho T_{max} - \frac{T_{max} - T}{\Delta}\right]}$ | (Lactin et al., 1995) | $\rho = 0.146 \pm\ 0.004$<br>$T_{max} = 38.5 \pm\ 0.3$<br>$\Delta = 6.82 \pm\ 0.19$<br>$AIC\ =\ -838.5$ |
| *lactin*2 | $R\ T\ = e^{\rho T} - e^{\left[\rho T_{max} - \frac{T_{max} - T}{\Delta}\right]} + \lambda$ | (Lactin et al., 1995) | $\rho = 0.103 \pm\ 0.011$<br>$\lambda = -0.074 \pm 0.026$<br>$T_{max} = 41.2 \pm\ 1.1$<br>$\Delta = 9.59 \pm\ 0.97$<br>$AIC\ =\ -856.41$ |
| *linear* | $R\ T\ = a + bT$ | (Campbell et al., 1974) | $a = -0.053 \pm \mathrm{XX}$<br>$\mathrm{b} = 0.007 \pm \mathrm{XX}$<br>$AIC\ =\ -780.06$ |
| *lrf* | $R\ T\ = r_{max} \cdot\ \tau\ ,$<br>where:<br>$\tau = \frac{T - T_{max}\ \cdot\ T - T_{min}\ ^2}{(T_{opt} - T_{min}) \cdot \left((T_{opt} - T_{min}) \cdot (T - T_{opt}) -\ T_{max}\ \cdot (T_{opt} + T_{min} - 2 \cdot T [illegible]}$ | (Rosso et al., 1993) | $r_{max} = 0.182 \pm 0.003$<br>$T_{opt} = 31.5 \pm 0.3$<br>$T_{min} = 3.97 \pm 1.21$<br>$T_{max} = 40.4 \pm 1.4$<br>$AIC\ =\ -857.35$ |
| *Mod_gaussian* | $R\ T\ =\ r_{max} \cdot e^{(-0.5(\frac{\lvert T - T_{opt}\rvert}{a})^b)}$ | (Lynch and Gabriel, 1987) | $r_{max} = 0.180 \pm 0.003$<br>$T_{opt} = 31.0 \pm 0.4$<br>$a = 9.63 \pm 5.52$<br>$b = 1.87 \pm 0.14$<br>$AIC = -857.53$ |
| *mod_polynomial* | $R\ T\ = a_0 + a_1 T + a_2 T^2 + a_3 T^3 + a_4 T^4$ | (Rebaudo et al., 2018) | Converged but was discarded (biologically unrealistic) |

| Model name | Equation | Reference | Parameter and AIC (estimate ± se; AIC) |
|---|---|---|---|
| *mod_weibull* | $R\ T = a\left(\frac{d-1}{d}\right)^{\frac{1-d}{d}}\left[\frac{T-T_{opt}}{c}+\left(\frac{d-1}{d}\right)^{\frac{1}{d}}\right]^{d-1} e^{-\left[\frac{T-T_{opt}}{c}+\left(\frac{d-1}{d}\right)^{\frac{1}{d}}\right]^{d}+\frac{d-1}{d}}$ | (Angilletta, 2006) | $a = 0.180 \pm 0.003$<br>$T_{opt} = 31.1 \pm 0.4$<br>$b = 39.6 \pm 8.5$<br>$c = 4.57 \pm 1.27$<br>$AIC = -857.64$ |
| *oneill* | $R\ T = r_{max}\cdot\left(\frac{CT_{max}-T}{CT_{max}-T_{opt}}\right)^{x}\cdot\ e^{x\cdot\frac{T-T_{opt}}{CT_{max}-T_{opt}}}$,<br>where: $x = \frac{w^2}{400}\cdot\left(1+\sqrt{1+\frac{40}{w}}\right)^2$,<br>and: $w = q_{10}-1 \cdot (CT_{max}-T_{opt})$ | (Krenek et al., 2011) | $r_{max} = 0.180 \pm 0.003$<br>$CT_{max} = 107 \pm 111$<br>$T_{opt} = 31.1 \pm 0.4$<br>$q_{10} = 1.95 \pm 0.07$<br>$AIC = -857.25$ |
| *pawar* | $R\ T = \frac{r_{Tref}\cdot e^{\frac{-E}{K}\left(\frac{1}{T+273.15}-\frac{1}{T_{ref}+273.15}\right)}}{1+\left(\frac{E}{E_H-E}\right)\cdot\ e^{\frac{E_H}{K}\left(\frac{1}{T_{opt}+273.15}-\frac{1}{T+273.15}\right)}}$ | (Kontopoulos et al., 2018) | $r_{Tref} = 0.250 \pm 0.009$<br>$e = 1.05 \pm 0.21$<br>$E_H = 1.30 \pm 0.09$<br>$T_{opt} = 35.2 \pm 3.0$<br>$AIC = -831.08$ |
| *schoolfield* | $R\ T = \frac{r_{Tref}\cdot e^{\frac{-E}{K}\left(\frac{1}{T+273.15}-\frac{1}{T_{ref}+273.15}\right)}}{1+e^{\frac{E_L}{K}\left(\frac{1}{T_L}-\frac{1}{T+273.15}\right)}+e^{\frac{E_H}{K}\left(\frac{1}{T_H}-\frac{1}{T+273.15}\right)}}$ | (Schoolfield et al., 1981) | $r_{Tref} = 0.169 \pm 0.022$<br>$e = 0.638 \pm 0.616$<br>$E_L = 1.81 \pm 1.63$<br>$T_L = 11.6 \pm 8.2$<br>$E_H = 1.89 \pm 0.54$<br>$T_H = 33.80 \pm 6.8$<br>$AIC = -854.09$ |

| Model name | Equation | Reference | Parameter and AIC (estimate ± se; AIC) |
|---|---|---|---|
| *thomas* | $R\ T\ = a \cdot e^{b \cdot T} \cdot (1 - (\frac{T - T_{ref}}{\frac{c}{2}})^2)$ | (Thomas et al., 2012) | $a = 0.024 \pm 0.005$<br>$b = 0.073 \pm 0.009$<br>$c = 33.2 \pm 1.1$<br>$T_{opt} = 23.7 \pm 0.9$<br>$AIC\ =\ -856.91$ |

### Appendix S4: Cold developmental thresholds

Here we define the *cold developmental thresholds* as the x-intercept values of the development rate models, which would be equivalent to the *lower development threshold* or *thermal base* typically described for linear, degree-day models and to the *critical thermal minimum* typically described for thermal performance curves of locomotive or survival rates. For the linear model, this value is calculated as $-\mathrm{a}/\mathrm{b}$ (see Table S1 above). For the nonlinear TPC models, while some of them incorporate this x-axis cutoff as an explicit parameter (see Table S1 above), we used the function *calc_params* from *rTPC* R package (Padfield et al., 2021) to systematize the method and work with average estimates. The resulting values of calculated cold developmental thresholds are given in Table S2.

**Table S2.** Cold developmental thresholds estimated for each fitted development model

| Development rate model | Cold developmental threshold (ºC) |
|---|---|
| *beta* | 7.29 |
| *boatman* | 7.58 |
| *briere1* | 7.22 |
| *briere2* | 6.76 |
| *flextpc* | 7.29 |
| *gaussian* | 6.57 |
| *joehnk* | 7.42 |
| *kamykowski* | 9.17 |
| *lactin1* | -0.721 |
| *lactin2* | 7.42 |
| *linear* | 7.12 |
| *lrf* | 8.46 |
| *mod_gaussian* | 5.86 |
| *oneill* | 5.71 |
| *pawar* | 2.83 |
| *schoolfield* | 7.44 |
| *thomas* | 7.04 |
| *weibull* | 6.68 |

### Appendix S5: Ranking of model validation

Table S3 and Table S4 show the calculated RMSEPs for each TPC model for daily and hourly temperature resolutions, respectively.

**Table S3**: RMSEP values (in days) for each fitted development rate model validated against averaged observations of CBMS transects when using daily temporatures from E-OBS at these sites.

| TPC model | RMSEP (days) |
|---|---|
| *lrf* | 7.49 |
| *briere2* | 9.01 |
| *briere1* | 9.58 |
| *schoolfield* | 9.64 |
| *boatman* | 9.69 |
| *beta* | 10.03 |
| *flextpc* | 10.03 |
| *thomas* | 10.31 |
| *linear_dd_rect* | 10.68 |
| *joehnk* | 10.90 |
| *lactin2* | 10.91 |
| *gaussian* | 11.44 |
| *weibull* | 11.80 |
| *mod_gaussian* | 15.43 |
| *oneill* | 15.89 |
| *kamykowski* | 16.08 |
| *linear_dd_triangle* | 16.62 |
| *linear_dd_sine* | 20.03 |
| *pawar* | 30.04 |
| *lactin1* | 40.42 |

**Table S4**: RMSEP values (in days) for each fitted development rate model validated against averaged observations of CBMS transects when using hourly temperatures downscaled from the originally daily temperatures from E-OBS at these sites.

| TPC model | RMSEP (days) |
|---|---|
| *kamykowski* | 6.34 |
| *joehnk* | 7.88 |
| *lactin2* | 7.88 |
| *thomas* | 8.20 |
| *briere1* | 9.29 |
| *briere2* | 10.19 |
| *boatman* | 12.07 |
| *linear_dd_rect* | 13.04 |
| *schoolfield* | 13.32 |
| *beta* | 13.57 |
| *flextpc* | 13.57 |
| *lrf* | 15.89 |
| *weibull* | 16.24 |
| *gaussian* | 16.30 |
| *mod_gaussian* | 19.84 |
| *oneill* | 20.28 |
| *pawar* | 31.86 |
| *lactin1* | 40.75 |

#### Appendix S6. Sensitivity analysis 1: fitting models separately to each development rate data set

Aiming to explore whether using disparate sources of developmental data across temperatures affects the outcome of the predictive accuracy exercise – likely due to both intrinsic differences in the source populations and to methodological disparities–, we repeated the analysis routine separately to each experimental data set (Gilbert, 1988; von Schmalensee et al., 2023).

Figures S5 and S6 show the validation outcome when including in the modelling pipeline only data from von Schmalensee et al., (2023) and from Gilbert, (1988), respectively.

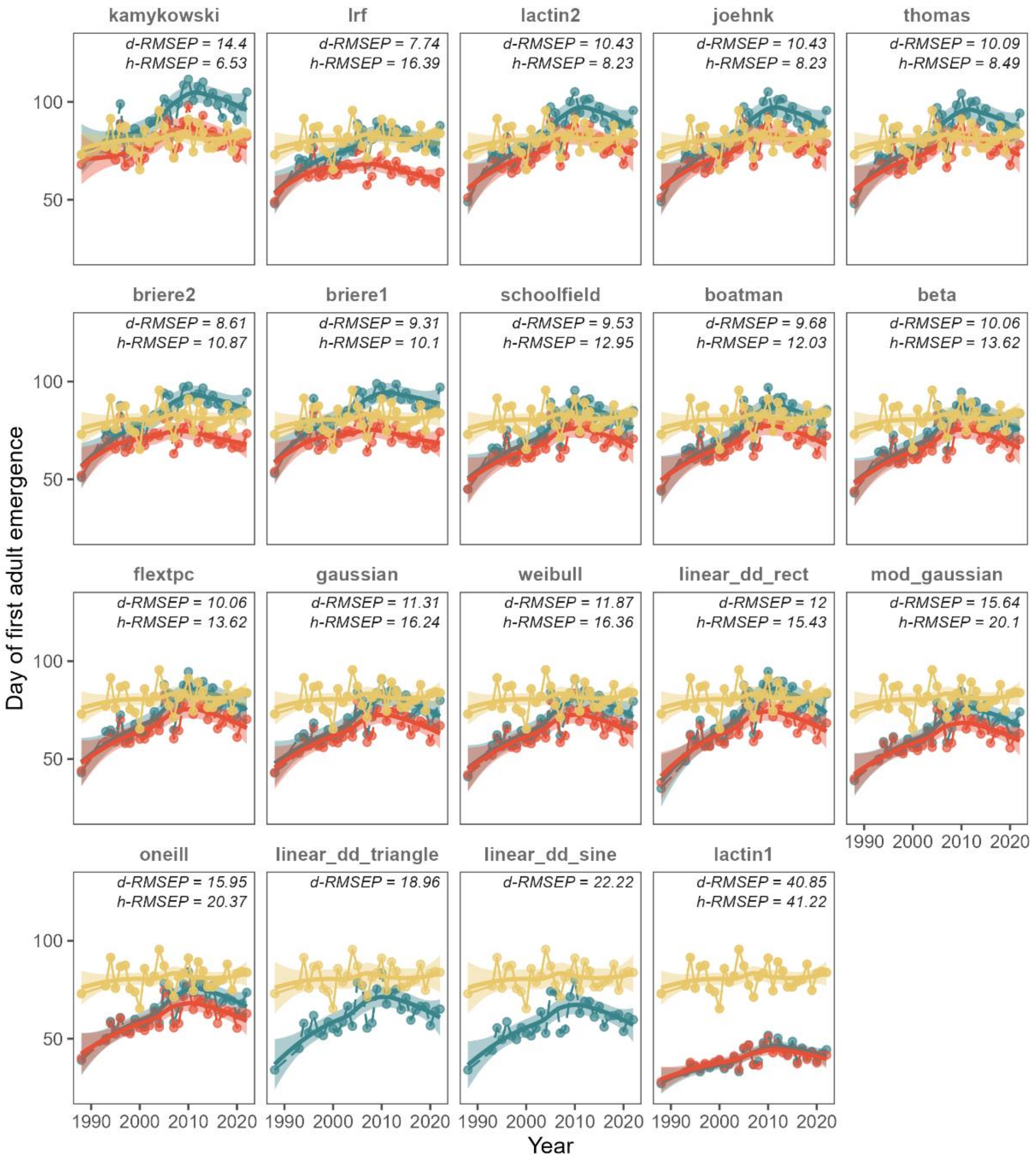


**Figure S5**: Validation of phenological predictions from model fitting using only data from von Schmalensee et al. (2023). Blue and red points represent phenological predictions from fitted models of first adult emergences, averaged across all transects, using daily and hourly temperature data, respectively. Predictions were accomplished by rate-summation for nonlinear TPC models and by degree-day calculations with rectangle, triangle and sine-methods for the linear model. Yellow points represent the averaged first observations of *P. rapae* adults across CBMS transects. Lines represent smooth trends with a loess model intended for facilitating visual comparison. RMSEPs are printed with abbreviations for both daily and hourly temperatures. Panels are ordered based on the lowest value of any RMSEP (either hourly or daily).

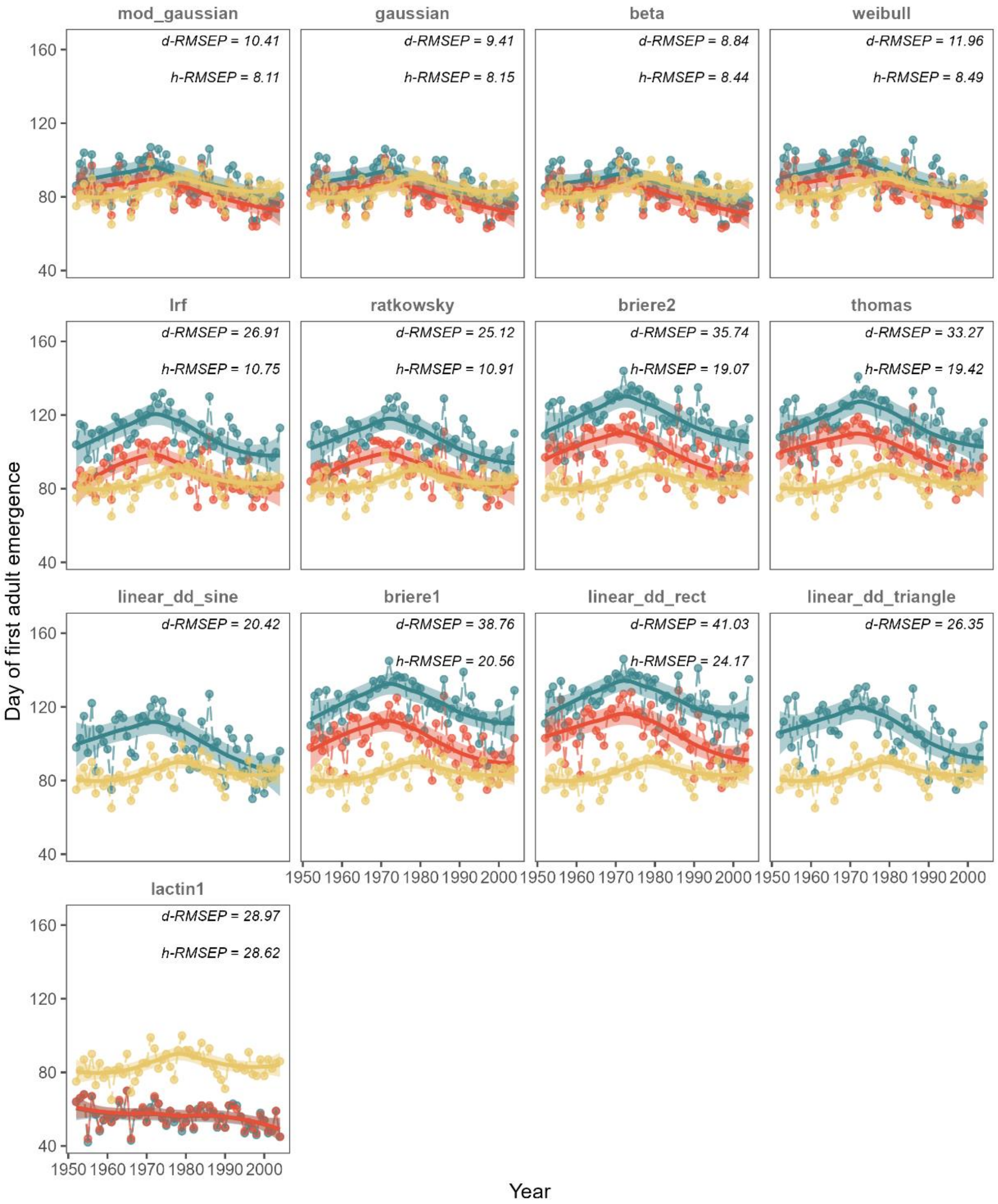


**Figure S6**: Validation of phenological predictions from model fitting using only data from Gilbert (1988). Blue and red points represent phenological predictions from fitted models of first adult emergences, averaged across all transects, using daily and hourly temperature data, respectively. Predictions were accomplished by rate-summation for nonlinear TPC models and by degree-day calculations with rectangle, triangle and sine-methods for the linear model. Yellow points represent the averaged first observations of *P. rapae* adults across CBMS transects. Lines represent smooth trends with a loess model intended for facilitating visual comparison. RMSEPs are printed with abbreviations for both daily and hourly temperatures. Panels are ordered based on the lowest value of any RMSEP (either hourly or daily).

#### Appendix S7. Validation using independent observational data of *P. rapae* first flights of the year from Gordo & Sanz (2006)

The validation of fitted model predictions against CBMS phenology data was repeated using an independent observational data set from (Gordo and Sanz, 2006). This data set compiles first emergences of P. rapae across mainland Spain for more than fifty years, and are available as averaged first emergence data points in their Figure 1. We obtained these values using WebPlotDigitizer v4.8 (https://apps.automeris.io/wpd4/).

Figure S7 shows this validation results.

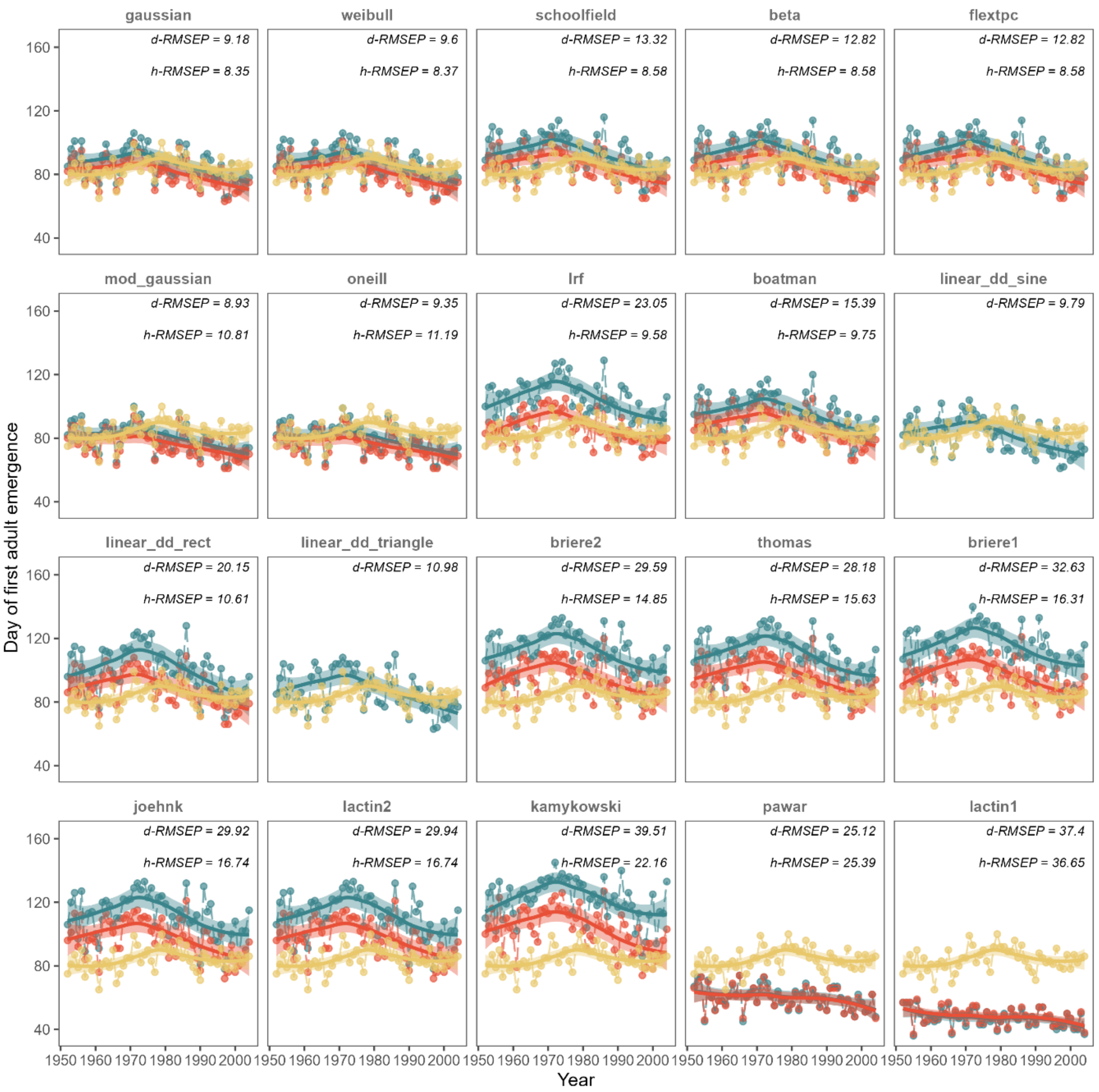


**Figure S7**: Validation of phenological predictions from model fitting to the combined development data sets against observational data from Gordo & Sanz (2006). Blue and red points represent phenological predictions from fitted models of first adult emergences, averaged across all transects, using daily and hourly temperature data, respectively. Predictions were accomplished by rate-summation for nonlinear TPC models and by degree-day calculations with rectangle, triangle and sine-methods for the linear model. Yellow points represent the averaged first observations of *P. rapae* adults mainland Spain, as reported in Figure 1 in Gordo & Sanz (2006). Lines represent smooth trends with a loess model intended for facilitating visual comparison. RMSEPs are printed with abbreviations for both daily and hourly temperatures. Panels are ordered based on the lowest value of any RMSEP (either hourly or daily).

### Appendix S8. CBMS data disaggregation for predicting phenology: the pooling effect

As explained in the main text, results shown in Figure 3 therein were obtained by calculating *RMSEPs* at both daily and hourly temperature resolutions and pooling all predicted and observed phenologies across transects. As discussed in the article, pooling and averaging phenologies lead to higher predictive accuracy. Here we show the equivalent results to Figure 2 when no pooling is carried out, i.e., when each observed first flight at each transect each year is compared against its corresponding prediction from rate summation approach to obtain a RMSEP and then these RMSEPs are averaged across transects and years. This alternative results in Figure S8, showing worsened predictive accuracy (i.e., larger distances between predictions in red/blue and observations in yellow), relative to the pooling approach presented in the main text.

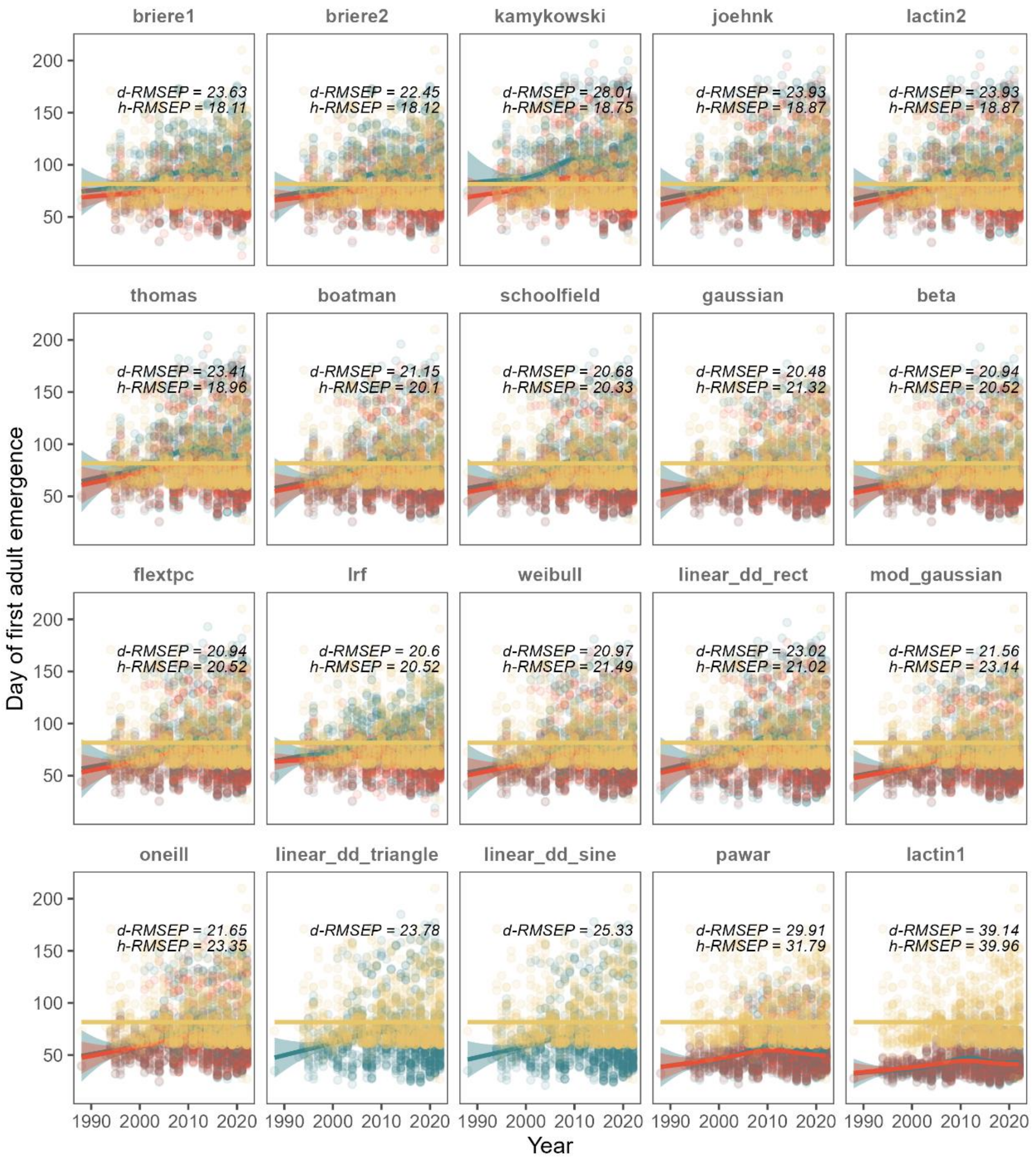


**Figure S8**: Validation of phenological predictions from TPC model fitting to combined Gilbert and Schmalensee datasets against disaggregated, observational data from the CBMS. Blue and red points represent phenological predictions from fitted models of first adult emergences using daily and hourly temperature data, respectively; each point represents predictions for a local CBMS transect. Predictions were accomplished by rate-summation (for nonlinear TPC models, i.e., all models except the linear) or by degree-day calculations (for linear). Yellow points represent the observed first adult flights of the year for each transect in the CBMS data series. Lines represent smooth trends with a loess model intended for facilitating visual comparison. RMSEPs represent are printed with abbreviations for both daily and hourly temperatures.

### Appendix S9. References in Supplementary Information:


Allen, J.C., 1976. A Modified Sine Wave Method for Calculating Degree Days 1. Environmental Entomology 5, 388–396. https://doi.org/10.1093/ee/5.3.388

Angilletta, M.J., 2006. Estimating and comparing thermal performance curves. Journal of Thermal Biology 31, 541–545. https://doi.org/10.1016/j.jtherbio.2006.06.002

Boatman, T.G., Lawson, T., Geider, R.J., 2017. A Key Marine Diazotroph in a Changing Ocean: The Interacting Effects of Temperature, CO2 and Light on the Growth of Trichodesmium erythraeum IMS101. PLoS One 12, e0168796. https://doi.org/10.1371/journal.pone.0168796

Briere, J.-F., Pracros, P., Le Roux, A.-Y., Pierre, J.-S., 1999. A Novel Rate Model of Temperature-Dependent Development for Arthropods. Environmental Entomology 28, 22–29. https://doi.org/10.1093/ee/28.1.22

Campbell, A., Frazer, B.D., Gilbert, N., Gutierrez, A.P., Mackauer, M., 1974. Temperature Requirements of Some Aphids and Their Parasites. Journal of Applied Ecology 11, 431–438. https://doi.org/10.2307/2402197

Chuine, I., Bonhomme, M., Legave, J.-M., García de Cortázar-Atauri, I., Charrier, G., Lacointe, A., Améglio, T., 2016. Can phenological models predict tree phenology accurately in the future? The unrevealed hurdle of endodormancy break. Global Change Biology 22, 3444–3460. https://doi.org/10.1111/gcb.13383

Cruz-Loya, M., Mordecai, E.A., Savage, V.M., 2024. A flexible model for thermal performance curves. bioRxiv 2024.08.01.605695. https://doi.org/10.1101/2024.08.01.605695

Curry, G.L., Feldman, R.M., Smith, K.C., 1978. A stochastic model of a temperature-dependent population. Theoretical Population Biology 13, 197–213. https://doi.org/10.1016/0040-5809(78)90042-4

Gilbert, N., 1988. Control of Fecundity in Pieris rapae. V. Comparisons Between Populations. Journal of Animal Ecology 57, 395–410. https://doi.org/10.2307/4913

Gordo, O., Sanz, J.J., 2006. Temporal trends in phenology of the honey bee Apis mellifera (L.) and the small white Pieris rapae (L.) in the Iberian Peninsula (1952–2004). Ecological Entomology 31, 261–268. https://doi.org/10.1111/j.1365-2311.2006.00787.x

Hernangómez, D., 2024. mapSpain: Administrative Boundaries of Spain.

Jöhnk, K.D., Huisman, J., Sharples, J., Sommeijer, B., Visser, P.M., Stroom, J.M., 2008. Summer heatwaves promote blooms of harmful cyanobacteria. Global Change Biology 14, 495–512. https://doi.org/10.1111/j.1365-2486.2007.01510.x

Kamykowski, D., 1986. A survey of protozoan laboratory temperature studies applied to marine dinoflagellate behavior from the viewpoint of ecological implications. (Appendix: Kamykowski, D. and F. G. Giesbrecht, Confidence limits for predicted swimming speed at a given temperature from S = SA (1-e-a(T-TL))(1-e-b(TH-T))). Presented at the Migration: Mechanisms and Adaptive Significance.

Kontopoulos, D.-G., García-Carreras, B., Sal, S., Smith, T.P., Pawar, S., 2018. Use and misuse of temperature normalisation in meta-analyses of thermal responses of biological traits. PeerJ 6, e4363. https://doi.org/10.7717/peerj.4363

Krenek, S., Berendonk, T.U., Petzoldt, T., 2011. Thermal performance curves of *Paramecium caudatum*: A model selection approach. European Journal of Protistology 47, 124–137. https://doi.org/10.1016/j.ejop.2010.12.001

Lactin, D.J., Holliday, N.J., Johnson, D.L., Craigen, R., 1995. Improved Rate Model of Temperature-Dependent Development by Arthropods. Environmental Entomology 24, 68–75. https://doi.org/10.1093/ee/24.1.68

Liu, S.-S., Zhang, G.-M., Zhu, J., 1995. Influence of Temperature Variations on Rate of Development in Insects: Analysis of Case Studies from Entomological Literature. Annals of the Entomological Society of America 88, 107–119. https://doi.org/10.1093/aesa/88.2.107

Luedeling, E., Caspersen, L., Fernandez, E., 2024. chillR: Statistical Methods for Phenology Analysis in Temperate Fruit Trees.

Lynch, M., Gabriel, W., 1987. Environmental Tolerance. The American Naturalist 129, 283–303. https://doi.org/10.1086/284635

Mirhosseini, M.A., Fathipour, Y., Reddy, G.V.P., 2017. Arthropod Development's Response to Temperature: a Review and New Software for Modeling. Annals of the Entomological Society of America 110, 507–520. https://doi.org/10.1093/aesa/sax071

Moore, J.L., Remais, J.V., 2014. Developmental Models for Estimating Ecological Responses to Environmental Variability: Structural, Parametric, and Experimental Issues. Acta Biotheor 62, 69–90. https://doi.org/10.1007/s10441-014-9209-9

Niehaus, A.C., Angilletta, M.J., Sears, M.W., Franklin, C.E., Wilson, R.S., 2012. Predicting the physiological performance of ectotherms in fluctuating thermal environments. J Exp Biol 215, 694–701. https://doi.org/10.1242/jeb.058032

Padfield, D., O'Sullivan, H., Pawar, S., 2021. rTPC and nls.multstart: A new pipeline to fit thermal performance curves in r. Methods in Ecology and Evolution 12, 1138–1143. https://doi.org/10.1111/2041-210X.13585

Rebaudo, F., Struelens, Q., Dangles, O., 2018. Modelling temperature-dependent development rate and phenology in arthropods: The devRate package for r. Methods in Ecology and Evolution 9, 1144–1150. https://doi.org/10.1111/2041-210X.12935

Rosso, L., Lobry, J.R., Flandrois, J.P., 1993. An unexpected correlation between cardinal temperatures of microbial growth highlighted by a new model. J Theor Biol 162, 447–463. https://doi.org/10.1006/jtbi.1993.1099

Sampaio, F., Krechemer, F.S., Marchioro, C.A., 2021. The hotter the better? Climate change and voltinism of *Spodoptera eridania* estimated with different methods. Journal of Thermal Biology 98, 102946. https://doi.org/10.1016/j.jtherbio.2021.102946

Schmalensee, L., Hulda Gunnarsdóttir, K., Näslund, J., Gotthard, K., Lehmann, P., 2021. Thermal performance under constant temperatures can accurately predict insect development times across naturally variable microclimates. Ecology Letters 24, 1633–1645. https://doi.org/10.1111/ele.13779

Schoolfield, R.M., Sharpe, P.J.H., Magnuson, C.E., 1981. Non-linear regression of biological temperature-dependent rate models based on absolute reaction-rate theory. Journal of Theoretical Biology 88, 719–731. https://doi.org/10.1016/0022-5193(81)90246-0

Stinner, R.E., Gutierrez, A.P., Butler, G.D., 1974. AN ALGORITHM FOR TEMPERATURE-DEPENDENT GROWTH RATE SIMULATION. Can Entomol 106, 519–524. https://doi.org/10.4039/Ent106519-5

Thomas, M.K., Kremer, C.T., Klausmeier, C.A., Litchman, E., 2012. A Global Pattern of Thermal Adaptation in Marine Phytoplankton. Science 338, 1085–1088. https://doi.org/10.1126/science.1224836

Vives-Ingla, M., Sala-Garcia, J., Stefanescu, C., Casadó-Tortosa, A., Garcia, M., Peñuelas, J., Carnicer, J., 2023. Interspecific differences in microhabitat use expose insects to contrasting thermal mortality. Ecological Monographs 93, e1561. https://doi.org/10.1002/ecm.1561

von Schmalensee, L., Caillault, P., Gunnarsdóttir, K.H., Gotthard, K., Lehmann, P., 2023. Seasonal specialization drives divergent population dynamics in two closely related butterflies. Nat Commun 14, 3663. https://doi.org/10.1038/s41467-023-39359-8
